# Phosphorylation of TTL3 by BIK1 Functions as a Molecular Switch to Control Cellulose Biosynthesis under Salt Stress

**DOI:** 10.64898/2026.06.25.734577

**Authors:** Francisco Percio, Raquel Pagano-Marquez, Angel Del Espino, Leia Colin, Jiamin Luo, Jessica Pérez-Sancho, Ryan Toth, Thomas A. DeFalco, Jian-Min Zhou, Alberto P. Macho, Cyril Zipfel, Lourdes Rubio, Staffan Persson, Vitor Amorim-Silva, Miguel A. Botella

**Author notes:** Authors for correspondence. **Contact information:** Amorim-Silva, Vitor, Botella, Miguel A. Equal contributions (Francisco Percio, Raquel Pagano-Marquez).

## Abstract

Cellulose, a central structural component of plant cell walls, is produced by cellulose synthase complexes (CSCs) at the plasma membrane. Salinity stress is particularly damaging to cellulose biosynthesis, and therefore, plants have developed adaptive mechanisms to cope with these conditions. TETRATRICOPEPTIDE THIOREDOXIN-LIKE (TTL) proteins are essential for growth under salt stress and show a salt-dependent association with CSCs through an as-yet unknown mechanism. Here, we identify a phosphorylation-dependent regulatory mechanism linking salt stress signaling to cellulose biosynthesis through the coordinated action of TTL3 and the receptor-like cytoplasmic kinase BOTRYTIS-INDUCED KINASE 1 (BIK1). Phosphorylation of Serine 93 in the N-terminal intrinsically disordered region of TTL3 controls its localization, retaining it in the cytosol, while dephosphorylation promotes association with CSCs at the plasma membrane. Biochemical and genetic analysis identified BIK1 as the kinase responsible for TTL3-S93 phosphorylation, with *bik1* mutants phenocopying the phosphoablative TTL3^S93A^ *in vivo*. Transcriptomic analyses reveal a strong overlap of differentially expressed genes between *bik1* and a cellulose-deficient mutant, supporting a broader role for BIK1 in cell wall regulation. Notably, TTL proteins do not appear to be involved in the assayed canonical immune responses, suggesting pathway specificity downstream of BIK1. Together, these findings define a signaling module that connects salt stress perception to CSCs regulation and establish BIK1-dependent TTL3 phosphorylation as a molecular switch for maintaining cell wall integrity under abiotic stress.

## Introduction

Soil salinity is one of the most serious environmental constraints worldwide, with detrimental effects on plant growth and crop production (Rosado et al., 2006; Shrivastava and Kumar, 2015; Zhang et al., 2022). The ability of the plant cell wall to remodel under high-salt conditions is considered a fundamental adaptation mechanism (Colin et al., 2023; Bawa et al., 2026). This is exemplified by the identification of *Arabidopsis thaliana* (hereafter Arabidopsis) mutants defective in cellulose biosynthesis by screening for salt hypersensitivity, such as *salt overly sensitive 5* (*sos5)* and *tetratricopeptide thioredoxin-like* (*ttl)* mutants (Shi et al., 2003; Rosado et al., 2006; Lakhssassi et al., 2012). In addition, mutants of genes encoding cellulose synthase complexes (CSCs) components, such as *companion of cellulose synthase* (*cc)*, *procuste 1* (*prc-1),* or *cellulose synthase interacting 1* (*csi1),* also show reduced tolerance to salt stress (Endler et al., 2015; Endler et al., 2016; Zhang et al., 2016). Cellulose microfibrils, the main load-bearing component of the cell wall, are directly synthesized at the plasma membrane by cellulose synthase (CESA) complexes; large protein complexes organized as six-fold symmetrical rosettes (Purushotham et al., 2020). In addition to the CESA catalytic core, other proteins/structures associated with CSCs contribute to cellulose synthesis, including cortical microtubules (MTs) that determine the direction of the synthesis by establishing the tracks of subcortical microtubules that guide the direction of cellulose synthesis in the plasma membrane (PM) (Pedersen et al., 2023). Proteins that are part of CSCs, i.e. move along the cellulose synthases, include (1) the CSI proteins that interacts with microtubules and CESAs to direct the synthesis of the microfibrils (Gu et al., 2010), (2) KORRIGAN, an endoglucanase that presumably trims or cleaves improperly formed cellulose segments (Vain et al., 2014), (3) CCs (Endler et al., 2015), and (4), TTL proteins (Kesten et al., 2022). The last two aid cellulose biosynthesis under salt stress. Moreover, in response to salt stress, MTs depolymerize, and CSCs get internalized into the cytosol in Small CESA Compartments (SmaCCs) (Endler et al., 2015). After the adaptation phase, with the collaboration of CCs and TTLs, MTs repolymerize in a stress-resilient pattern, and CSCs are recovered to the PM (Endler et al., 2015; Kesten et al., 2022). As a result, cellulose biosynthesis is restored, generating a new array of cellulose microfibrils that contributes to adaptation to high salinity, thereby allowing further cellular growth (Endler et al., 2015). A distinctive feature of TTL proteins compared with other CSC-associated components is their dynamic recruitment to cellulose synthase complexes. Under normal conditions, TTLs are predominantly cytosolic, whereas salt stress or genetic perturbations that reduce cellulose levels promote their association with CSCs (Kesten et al., 2022).

TTL proteins exhibit a common modular architecture containing an intrinsic disordered region at the N-terminus (N-IDR), six tetratricopeptide repeat (TPR) domains distributed in specific positions throughout the sequence, and a C-terminal sequence with homology to thioredoxins (Rosado et al., 2006; Lakhssassi et al., 2012). Intrinsically disordered regions (IDRs) are often regulatory targets for post-translational modifications, which, due to their structural flexibility, undergo rapid conformational changes that modulate protein activity, interactions, and cellular localization (Iakoucheva et al., 2004; Diella et al., 2008). TPR domains are widespread protein-protein interaction domains involved in multiple cellular processes, with partners ranging from short peptides to large proteins (Perez-Riba and Itzhaki, 2019).

The receptor-like cytoplasmic kinase BOTRYTIS-INDUCED KINASE 1 (BIK1) is a central regulator of plant immunity, transducing signals from multiple pattern recognition receptors to activate downstream defenses (Lu et al., 2010; Dodds et al., 2024; Hailemariam et al., 2024). Interestingly, BIK1 phosphorylates SHOU4 and SHOU4L; proteins involved in CSCs trafficking (Wang et al., 2023), hinting at a connection between immune signaling and cellulose biosynthesis. However, direct links between BIK1 and CESA-associated proteins under abiotic stress conditions remain elusive.

Here, we show that phosphorylation of Ser93 in the TTL3 N-IDR serves as a molecular switch that controls TTL3 association with the CSCs. Furthermore, we identify BIK1 as the kinase responsible for directly phosphorylating TTL3 at Ser93. Notably, while BIK1 controls TTL3 phosphorylation and localization, TTL proteins do not contribute to the canonical immune responses tested here, supporting pathway-specific functions downstream of this multifunctional kinase.

## Results

### TTL3 is phosphorylated at the N-terminal intrinsically disordered region

Under control conditions, TTL3 is mainly localized in the cytosol, but a fraction also engages with the CSCs through interaction with CESA1 (Kesten et al., 2022). Conditions that reduce cellulose, such as salt stress or genetic manipulation, trigger substantial relocalization of TTL3 from the cytosol to PM-localized CSCs, leading to CSCs stabilization (Kesten et al., 2022). To investigate the mechanisms underlying TTL3 localization dynamics, we first analyzed whether TTL3 carries post-translational modifications (PTMs) using a functional TTL3-GFP Arabidopsis line (Amorim-Silva et al., 2019) and immunoprecipitation-mass spectrometry (IP-MS). PTMs, and prominently phosphorylation, are commonly used to regulate protein activity, interactions, and cellular dynamics. Interestingly, we found that TTL3 may be phosphorylated at residues S8, S42, S93, S98, T184, S235, and S249 (Fig. S1A, Fig. S2). Here, the first five residues are part of the N-terminal intrinsically disordered region (N-IDR) of TTL3 (Fig. 1A). We corroborated the first four phosphorylated residues through the Arabidopsis phosphoproteomic database (https://www.psb.ugent.be/webtools/ptm-viewer/) (Fig. S1A).

**Fig. 1.**
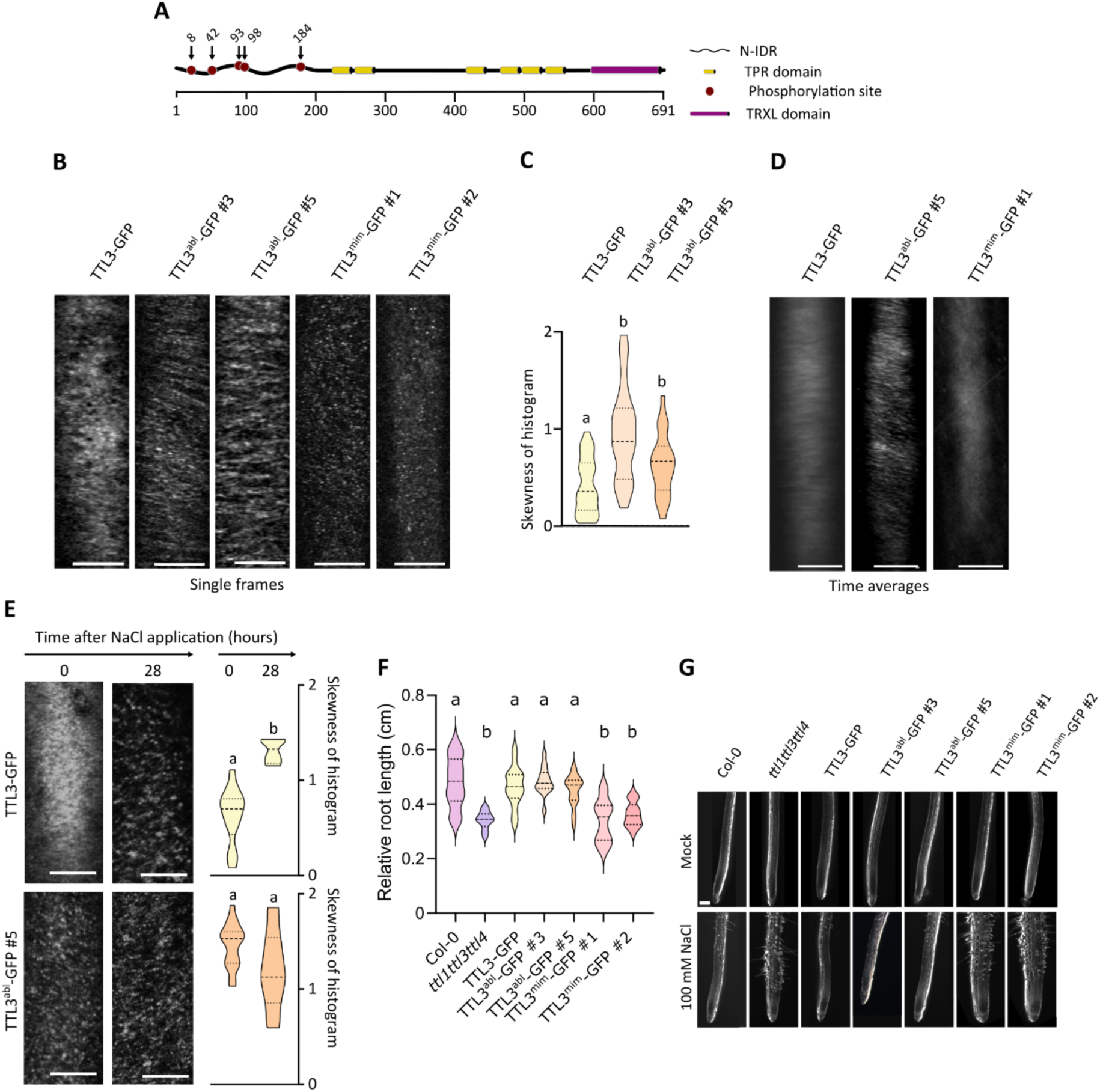
TTL3 phosphorylation controls its localization and function in cellulose biosynthesis. **A.** TTL3 protein model. Domains were predicted using InterPro (https://www.ebi.ac.uk/interpro/) and phosphorylation sites using our IP-MS data and data from Mergner et al. (2020). **B.** Representative single frame spinning disk confocal microscopy images of 2.5-days-old etiolated hypocotyl epidermal cells of *ttl1ttl3ttl4* TTL3-GFP and two independent lines of TTL3^abl^-GFP and TTL3^mim^-GFP. Scale bar 5 μm. **C.** Skewness of histogram of (B). One-way ANOVA analysis with Tukey’s multiple comparative test, P≤0.05, N>20 independent cells from at least 9 different seedlings for each line. Repeated twice with similar results. **D.** Two-minutes time average images for TTL-GFP and its mutated versions. Scale bar 5 μm. **E.** Spinning disk confocal microscopy imaging of 2.5-days-old etiolated hypocotyl epidermal cells of *ttl1ttl3ttl4* TTL3-GFP and TTL3^abl^-GFP 5.10 exposed to 200 mM NaCl. Images represent single frames taken at the indicated times of exposure. Time 0 shows cells just before NaCl treatment. One-way ANOVA analysis with Tukey’s multiple comparative test, P≤0.05, N>10 independent cells from at least 4 different seedlings for each line. Repeated twice with similar results. Scale bar 5 μm. **F.** WT (Col-0) and *ttl1ttl3ttl4* mutants and two TTL3^abl^-GFP and TTL3^mim^-GFP independent lines were germinated and grown for 3 days under control conditions and then transferred to control media or media supplemented with 100 mM NaCl and grown for seven additional days. N ≥ 20 roots and three independent experiments with similar results. One way-ANOVA with Tukey’s multiple comparative test and p<0.001. **G.** Magnifications of root tip from (F). Scale bar 250 μm.

### TTL3 phosphorylation regulates its association with CSCs

We next aimed to determine whether the N-IDR-associated phosphosites affected previously described cytosol-PM dual localization of TTL3 (Kesten et al., 2022). For this, we generated two different versions of *pTTL3:TTL3-GFP*, a phosphoablative version in which the four serine (S) and threonine (T) residues were changed to alanine (A) to produce *pTTL3:TTL3^S8,S42,S93,S98,T184/A^-GFP* (hereafter TTL3^abl^-GFP); and a phosphomimetic version in which the five residues were changed to aspartic acid (D) *pTTL3:TTL3^S8,S42,S93,S98,T184/D^* (hereafter TTL3^mim^-GFP). We transformed these constructs into the triple Arabidopsis mutant *ttl1ttl3ttl4* and selected two independent homozygous lines based on GFP fluorescence of the phosphoablative (#3 and #5) and the phosphomimetic (#1 and #2) lines for further analysis.

To investigate if N-IDR phosphorylation status changes TTL3 localization dynamics, we imaged four-day-old etiolated hypocotyls using spinning-disk confocal microscopy. Focusing on the PM, we followed established protocols (Verbančič et al., 2021) with minor modifications to improve handling and sample fitness (see Methods). As reported (Kesten et al., 2022), TTL3-GFP showed a prominent cytosolic localization, with some linear tracks resulting from its association with the CSCs at the PM (Fig. 1B). Interestingly, TTL3*^abl^*-GFP lines displayed an apparent increase in CSCs to cytosol ratio, suggesting a reduced cytosolic signal and increased CSCs association (Fig. 1B). Analysis of TTL3 clustering at the PM was carried out through the quantification of the skewness of the histogram of several images as previously described (Kesten et al., 2022). TTL3-GFP exhibited lower values of skewness, consistent with its mainly-cytosolic diffuse localization, while TTL3^abl^-GFP exhibited higher values of skewness due to its clustering at the PM, along with the CSCs (Fig. 1C) (Kesten et al., 2022). This data supports the hypothesis that the lack of phosphorylation in these residues enhanced TTL3 localization to the CSCs. Moreover, time-lapse imaging showed that migratory TTL3-GFP fluorescence was substantially obscured by strong cytosolic signals, whereas TTL3^abl^-GFP mostly showed tracks resembling CSCs dynamics (Fig. 1D), supporting the hypothesis that the inability to phosphorylate these N-IDR residues enhances TTL3 localization to the CSCs. The TTL3^mim^-GFP lines showed a punctuated signal, neither similar to the TTL3-GFP cytosolic signal nor to the PM localization one (Fig. 1B). Time-lapse images of TTL3^mim^-GFP showed a diffuse pattern (Fig. 1D), suggesting that the fluorescent puncta observed in single time point images were likely the result of protein aggregations that localized across PM and/or the cytosol that do not follow the CSCs. Hence, we did not calculate the skewness in these lines, since although TTL3^mim^-GFP was aggregated, it did not migrate with CSCs as TTL3^abl^-GFP does.

Next, we investigated the role of N-IDR phosphorylation in the localization and dynamics of TTL3 during salt stress, focusing on TTL3-GFP and TTL3^abl^-GFP #5. We analyzed control conditions (0 hour) and 28 hours after treatment with 200 mM NaCl. TTL3-GFP was mostly cytosolic at 0 hour and was recruited to the CSCs after 28 hours of NaCl treatment (Fig. 1E; Kesten et al., 2022). By contrast, in TTL3^abl^-GFP lines, NaCl treatment did not cause obvious changes in the distribution between cytosol and PM (Fig. 1E). In summary, our data indicate that the lack of phosphorylation at these five N-IDR residues constitutively targets TTL3^abl^-GFP to the PM, enabling it to bind the CSCs independently of stress conditions.

Root growth analyses of these lines indicated that, under control conditions, *ttl1ttl3ttl4* shows root length and anisotropic growth similar to WT (Col-0) plants, while under salt stress, *ttl1ttl3ttl4* shows reduced root growth and isotropic root cell expansion, defects that were complemented by TTL3-GFP (Figs. 1F, G, and S1B). Interestingly, TTL3^abl^-GFP lines complemented the phenotypes of the *ttl1ttl3ttl4* mutant, whereas TTL3^mim^-GFP lines did not (Figs. 1F, G, and S1B). Taken together, these results indicated that while the phosphoablative TTL3 may rescue the salt-defective growth of *ttl1ttl3ttl4*, the phosphomimetic TTL3 constructs failed to do so.

### TTL3 dynamics and function are controlled by Serine 93 phosphorylation

We next aimed to determine whether individual phosphosites of the N-IDR were responsible for TTL3 localization and dynamics. To this end, we first established a protocol to analyze the localization of TTL3-GFP, TTL3^abl^-GFP, and TTL3^mim^-GFP using transient expression in *Nicotiana benthamiana,* and imaging their localization patterns at the cortical plane of leaf cells by confocal microscopy. TTL3-GFP expressed in *N. benthamiana* leaves showed dual localization, similar to that in a stably transformed Arabidopsis line under control conditions: a diffuse pattern indicating cytosolic localization and straight lines consistent with CSC-localized TTL3 (Fig. S3A). TTL3^abl^-GFP expressed in *N. benthamiana* mostly showed signals resembling CSCs tracks, while TTL3^mim^-GFP mostly presented a diffuse signal, in both cases resembling Arabidopsis localization (Fig. S3A). These results support the use of *N. benthamiana* as a proxy to analyze the subcellular localization of different TTL3 versions. We next generated *pTTL3:TTL3-GFP* constructs in which S8, S42, S93, S98, and T184 were individually mutated to alanine (Fig. S3B). Each construct was expressed in *N. benthamiana,* and the localization of each TTL3 mutant was analyzed (Fig. S3C). TTL3^S8A^-GFP, TTL3^S42A^-GFP, TTL3^S98A^-GFP, and TTL3^T184A^-GFP showed patterns reminiscent of TTL3-GFP. However, TTL3^S93A^-GFP showed a pattern mainly resembling that of TTL3^abl^-GFP, suggesting that S93 of TTL3 is responsible for its targeting to the PM.

To further investigate the role of S93 in TTL3 dynamics, we generated stable Arabidopsis *ttl1ttl3ttl4* lines expressing *pTTL3:TTL3^S93A^GFP* (see Methods). Confocal microscopy analysis of two independent *pTTL3:TTL3^S93A^GFP* lines (lines #9 and #14) showed a pattern similar to that of *TTL3^abl^-GFP* lines with a main localization at PM-CSC tracks (Fig. 2A). This result indicates that the phosphorylation status of S93 is responsible for TTL3 localization with the CSCs, and that the lack of phosphorylation in this residue guides its interaction with CSCs at the PM.

**Fig. 2.**
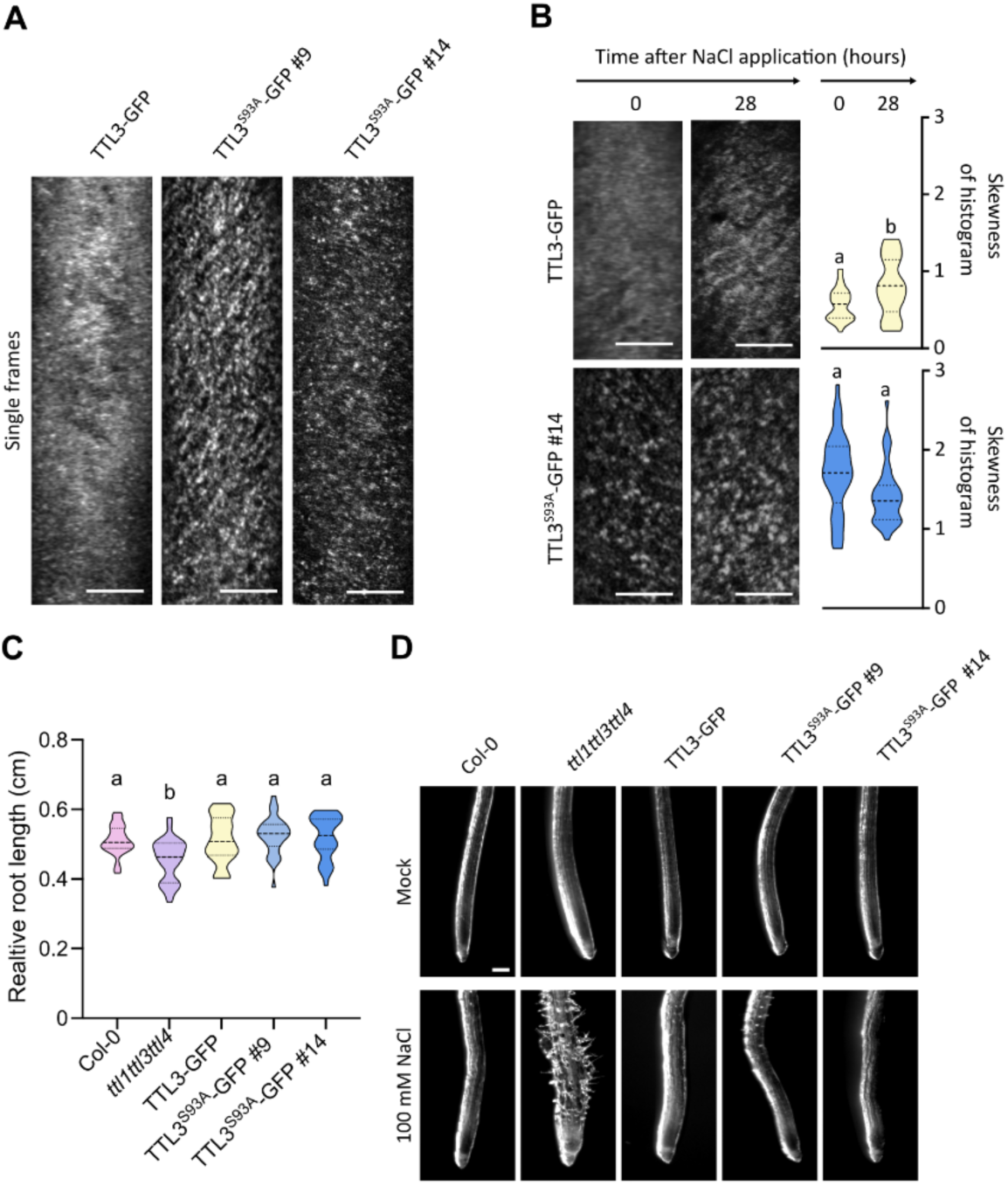
Phosphorylation status of TTL3-S93 controls TTL3 localization and salt stress tolerance of Arabidopsis seedlings. **A.** Representative single frame spinning disk confocal microscopy images of 2.5-days-old etiolated hypocotyl epidermal cells of *ttl1ttl3ttl4* TTL3-GFP and two independent lines of TTL3^S93A^-GFP. Scale bar 5 μm. **B.** Confocal microscopy imaging of 2.5-days-old etiolated hypocotyl epidermal cells of *ttl1ttl3ttl4* TTL3-GFP and TTL3^S93A^-GFP 14.1 exposed to 200 mM NaCl. Images represent single frames taken at the indicated times of exposure. Time 0 shows cells just before NaCl treatment. One-way ANOVA analysis with Tukey’s multiple comparative test, P≤0.05, N>30 independent cells from at least 9 different seedlings for each line, three independent replicates. Scale bar 5 μm. **C.** WT (Col-0) and *ttl1tt3ttl4* mutants and two TTL3^S93A^-GFP independent lines were germinated and grown for 3 days under control conditions and then transferred to control media or media supplemented with 100 mM NaCl and grown for seven additional days. N ≥ 20 roots and three independent experiments. One way-ANOVA with Tukey’s multiple comparative test and p<0.001, three independent replicates with similar results. **D.** Magnifications of root tip from (C). Scale bar 250 μm.

We then analyzed the localization and dynamics of TTL3^S93A^-GFP after NaCl treatment. Like TTL3^abl^-GFP, TTL3^S93A^-GFP showed a CSC-like dotted and linear localization in control conditions, which was maintained after 28 hours of NaCl treatment, indicating that the lack of S93 phosphorylation caused constitutive TTL3 targeting to the PM CSCs independently of salt stress (Fig. 2B). We further determined the role of TTL3-S93 on root length (Figs. 2C and S4) and anisotropic growth (Fig 2D) under salt stress. TTL3^S93A^-GFP complemented the growth defects of the *ttl1ttl3ttl4* mutant as TTL3^abl^-GFP did, indicating that S93 is the key residue controlling TTL3 localization, dynamics, and function.

### BIK1 phosphorylates TTL3 at serine 93

Kinase substrate recognition is regulated by multiple factors, including short linear motifs immediately proximal to targeted phosphorylation sites. The S93 site within the TTL3 N-IDR of TTL3 conforms to the recently described substrate motif of BIK1 (Toth et al., 2026), suggesting that this site could be targeted by this kinase. Notably, this putative BIK1 motif sequence was identified in the N-IDR of TTL proteins in Arabidopsis and other species, including distant species such as *Marchantia polymorpha* (Fig. 3A, B), highlighting the potential importance of this putative phosphorylation site.

**Fig. 3.**
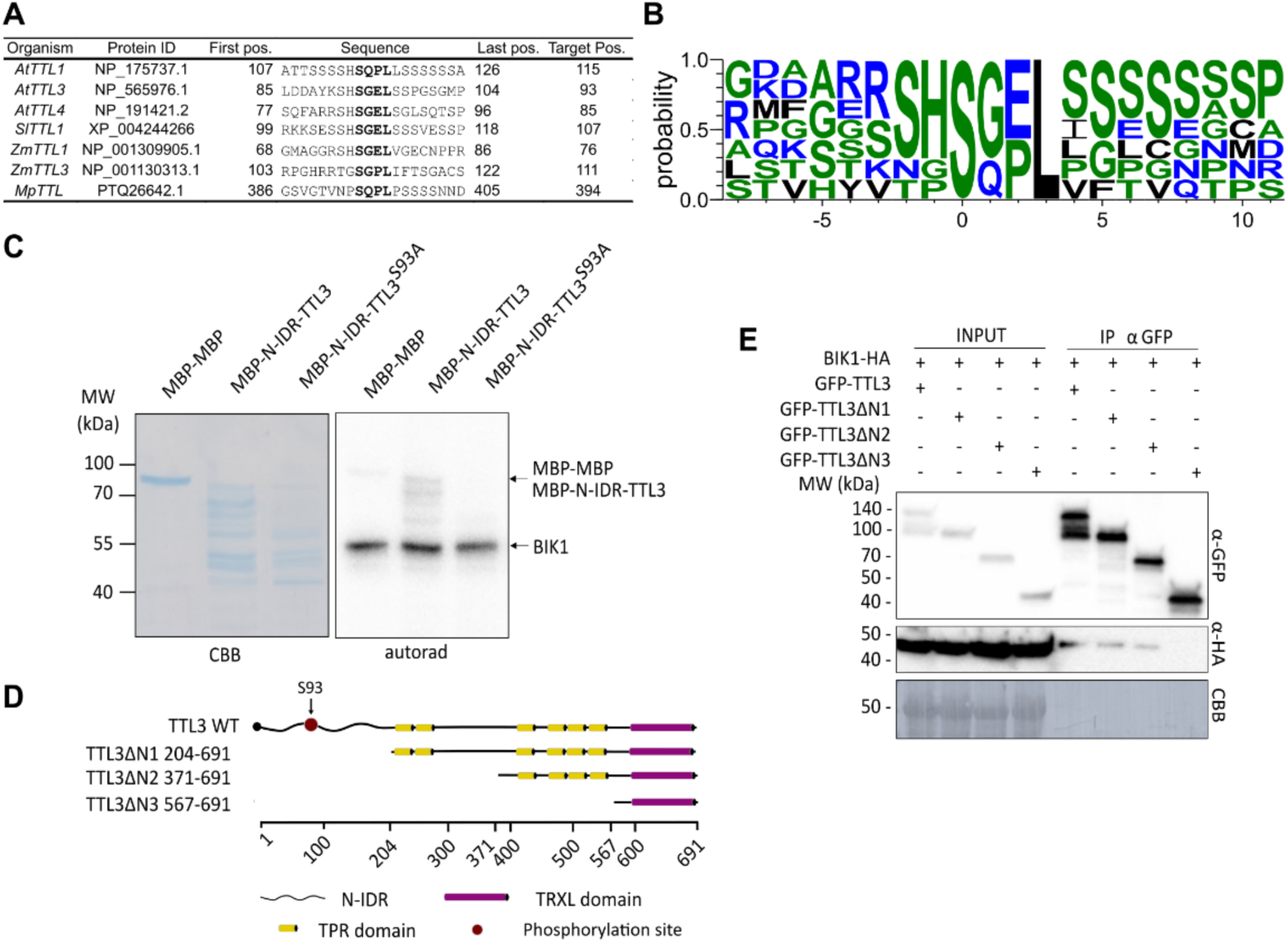
BIK1 phosphorylates AtTTL3 and other TTLs across multiple species. **A.** Predicted target sequence for BIK1 phosphorylation in TTLs of *Arabidopsis thaliana*, *Solanum lycopersicum*, *Zea mays* and *Marchantia polymorpha*. Target sequence is highlighted, the first serine being the one predicted to be phosphorylated. **B.** Consensus sequence for BIK1 phosphorylation in the proteins depicted at (A) (made with https://weblogo.threeplusone.com/). **C.** Transphosphorylation assay of TTL3 by BIK1. N-terminus of TTL3 was expressed in *E. coli* and purified using MALTOSE-BINDING PROTEIN (MBP). MBP-MBP was used as a negative control, and a truncated version of TTL3 in which the serine 93 was substituted for alanine was used to check phosphorylation specificity. CBB: Coomassie Brilliant Blue. **D.** Schematic representation of TTL3 WT and truncated versions used in (E). **E.** GFP-TTL3 and N-terminal truncated versions of TTL3 fused to eGFP were co-expressed in *Nicotiana benthamiana* with BIK1-HA_3_. Input and Co-IP proteins using α-GFP beads were analyzed by immunoblot using anti-GFP and anti-HA antibodies. Equal loading was confirmed by Coomassie Brilliant Blue (CBB) staining of input samples.

The ability of BIK1 to directly phosphorylate TTL3 was therefore investigated using an *in vitro* kinase assay. Due to the instability of the complete TTL3 protein, the N-IDR of TTL3 was fused to MALTOSE BINDING PROTEIN (MBP) (MBP-TTL3) and used for affinity purification. A tandem MBP dimer was used as a negative control, and a MBP-TTL3^S93A^ mutant was used to determine the specificity of the phosphorylation. BIK1 was able to directly phosphorylate the WT MBP-TTL3 fusion but not MBP-TTL3^S93A^ (Fig. 3C), indicating that BIK1 specifically targets this site *in vitro*.

Next, we showed that TLL3 associates *in vivo* with BIK1 using coimmunoprecipitation (Co-IP) assays in *N. benthamiana* (Fig. 3D, E). To define the TTL3 domain that associates with BIK1, we included several truncated TTL3 constructs and performed Co-IP with BIK1. The constructs comprised TTL3 without the N-IDR (TTL3ΔN1), TTL3 without the N-IDR and the first two TPRs (TTL3ΔN2), and a construct comprising only the TRXL domain (TTL3ΔN3) (Fig. 3D). As shown in Fig. 3E, TTL3ΔN1 and TTL3ΔN2 but not TTL3ΔN3 co-immunoprecipitated BIK1 to a similar extent as the full-length TTL3, suggesting that the last four TPRs are the domains required for TTL3 association with BIK1.

### BIK1 regulates TTL3 localization via Ser 93 phosphorylation

If BIK1 is the kinase responsible for the S93 phosphorylation *in vivo*, we hypothesized that in an Arabidopsis *bik1* mutant background, TTL3-GFP would show the same localization as TTL3^S93A^-GFP. To investigate this, we generated *BIK1* loss-of-function mutants in the *ttl1ttl3ttl4 pTTL3:TTL3-GFP* background using CRISPR/Cas9 (see Methods). These new alleles were named *bik1-11* and *bik1-12* based on the *bik1* alleles previously generated (Song et al., 2026). The BIK1 proteins encoded in the TTL3-GFP *bik1-11* and TTL3-GFP *bik1-12* lines are 90- and 82-amino acids long, respectively, and are predicted to be loss-of-function mutants since they lack most of the kinase catalytic domain of BIK1 (Fig. 4A). Interestingly, like TTL3^S93A^-GFP, both *bik1* mutant alleles lost the diffuse TTL3-GFP cytosolic signal (Fig. 4B), suggesting a further association of TTL3-GFP to the CSCs at the PM. Next, we quantified the total amount of TTL3-GFP in these lines compared to the original TTL3-GFP line using immunoblots. Strikingly, loss of *BIK1* function caused a drastic reduction of TTL3-GFP of about 80 % (Fig. 4C). As expected, *BIK1* loss-of-function mutant alleles such as *bik1-2* and *bik1-3* (Song et al., 2026) show WT responses to NaCl, phenocopying TTL3^S93A^-GFP (Figs. 4D, 4E, and S5), consistent with BIK1 role of phosphorylating S93.

**Fig. 4.**
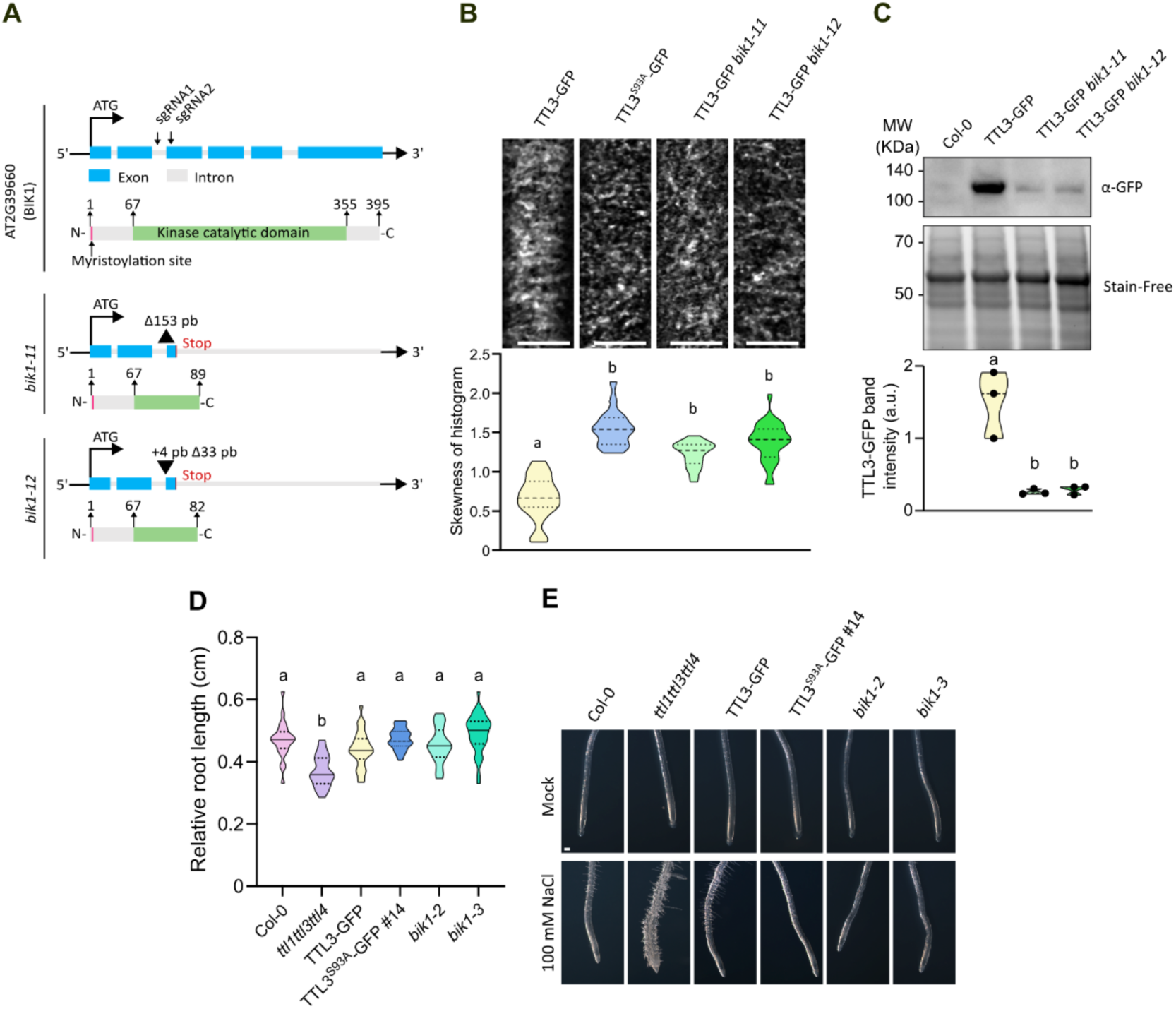
Lack of BIK1 alters TTL3 localization. **A.** Schematic representation of novel generated CRISPR/Cas-9 *bik1* mutant lines. Genomic and protein sequence of AT2G39660 (BIK1) are depicted, with exon and introns or the relevant protein domains specified respectively. **B.** Representative single frame confocal microscopy images of 2.5-days-old etiolated hypocotyl epidermal cells of *ttl1ttl3ttl4* TTL3-GFP and two independent lines of CRISPR/Cas-9 BIK1 mutated TTL3-GFP (top panel) and its quantified skewness (bottom panel). One way-ANOVA with Tukey’s multiple comparative test, P ≤ 0.05, three independent replicates, N>25 independent cells from at least 9 different seedlings for each line. Scale bar 5 μm. **C.** Inmunoblot analysis of TTL3-GFP in WT and two independent *bik1* loss-of-function mutant backgrounds, immunoblot using anti-GFP to detect TTL3-GFP and Stain free (BioRad) of the total protein to confirm equal loading. **E.** WT (Col-0), *ttl* and *bik1* mutants and two independent *bik1* loss-of-function mutant lines were germinated and grown for 3 days under control conditions and then transferred to control media or media supplemented with 100 mM NaCl and grown for seven additional days. N ≥ 20 roots and One way-ANOVA with Tukey’s multiple comparative test and p<0.001, three independent replicates with similar results. **F.** Magnifications of root tip from (E). Scale bar 250 μm.

### *bik1* and the cellulose-deficient mutant *prc1-1* show similar transcriptional changes

To further investigate the possible role of BIK1 in cellulose biosynthesis, we analyzed changes in the transcriptome of the *bik1-2* mutant (Song et al., 2026) by RNA-seq and compared it with that of the cellulose synthase (*CESA6*) mutant *prc1-1* (Fagard et al., 2000). The analysis (listed in Table S1, S2, S3) showed 921 differentially expressed genes (DEG) between *bik1-2* and WT (Fig. 5A), 660 up and 261 down (Fig. 5B), and 1,594 DEG between *prc1-1* and WT (Fig. 5A), 1,065 up and 529 down (Fig. 5B). Remarkably, *bik1-2* showed around 80 % of common DEGs with *prc1-1* (Fig. 5A) with similar expression trends (Fig. 5B, C) (see Methods). This unexpected representation factor showed that the gene sets overlapped 13.2 times greater compared with same-sized independent gene sets, and the probability that this overlap occur by chance is extremely close to zero (p<0.000e+00). The analysis of sample grouping in the principal component analysis (PCA) plot and the profiles of their volcano plots supported the conclusion that *bik1-2* and *prc1-1* have similar transcriptional programs (Fig. S6). Of note, the PCA analysis showed that PC1 (57.32% of the variance) clearly separates the WT group from the *bik1-2* and *prc1-1* mutant groups. This means that the greatest biological difference in transcriptional data lies between the WT and the mutants. Transcriptome assembly allowed us to confirm the previously described mutations in the *bik1-2* and *prc1-1* mutant lines used in this analysis (Fig. S6). Functional enrichment analysis of overlapping DEGs for *bik1-2* and *prc1-1* revealed the highest overrepresentation of GO terms related to abiotic and biotic stress responses (Fig. 5D). The large overlap of DEGs between *bik1-2* and *prc1-1* further supports the role of *BIK1* in cellulose biosynthesis.

**Fig. 5.**
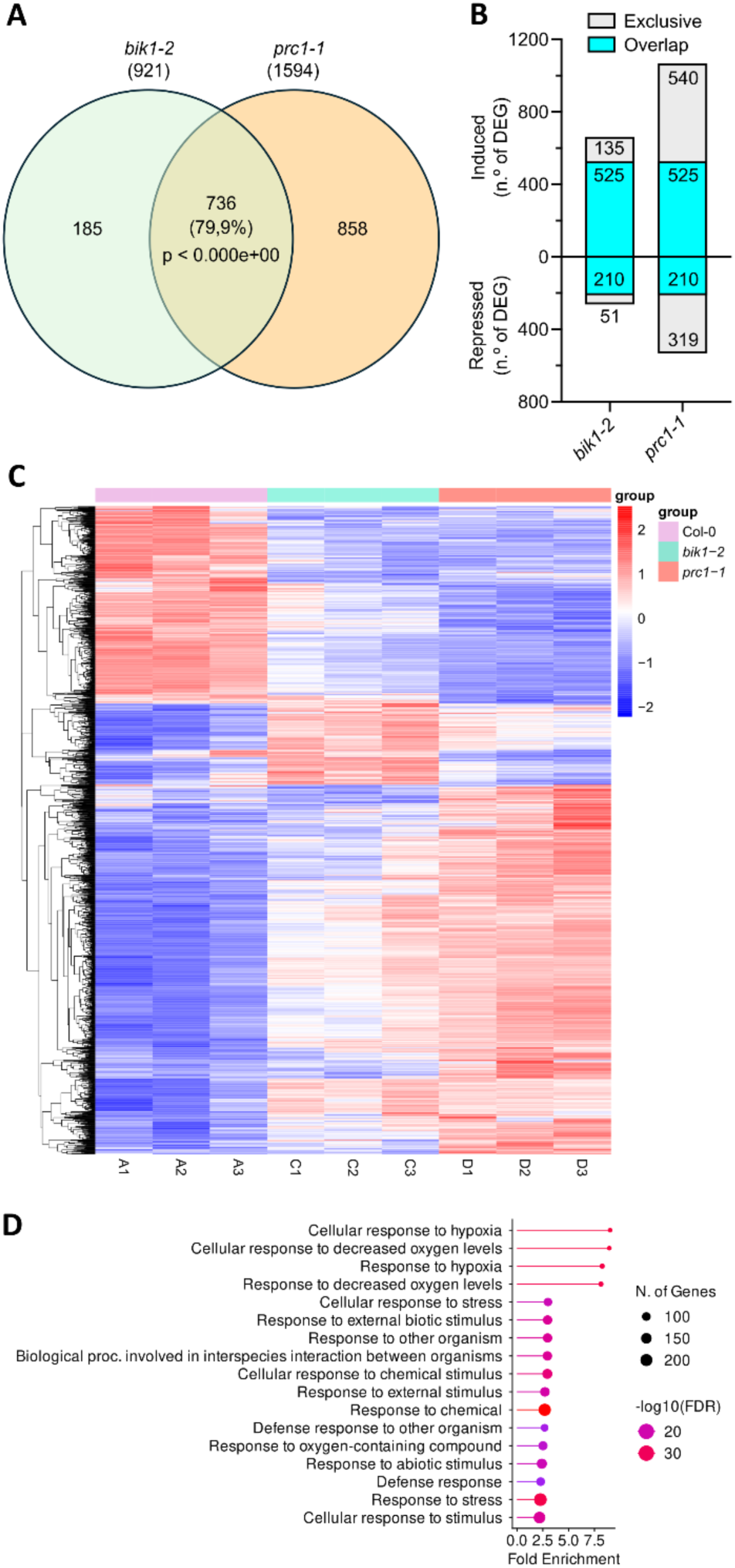
RNAseq analysis in rosettes of Arabidopsis plants with four-weeks old. **A.** Overlap of *bik1-2* and *prc1-1* DEG with induced and repressed DEG analyzed together. **B.** Overlap of *bik1-2* and *prc1-1* DEG with induced and repressed DEG analyzed independently. **C.** Heat-map with the Z-score normalization of the expression data (FPKM) for WT *bik1-2* and *prc1-1.* **D.** Gene ontology biological process enriched categories for in DEG that overlap between *bik1-2* and *prc1-1*.

### TTL proteins do not play a role in tested canonical immune responses

BIK1 is a key component of plant immune responses to multiple pathogens, including *Pseudomonas syringae* (Lu et al., 2010; Zhang et al., 2010; Lei et al., 2014). Moreover, SHOU4 and the related SHOU4L, with a reported role in trafficking of the CSCs to the PM (Polko et al., 2018), are phosphorylated by BIK1, contributing to plant immunity (Wang et al., 2023). Therefore, we investigated whether *TTL* genes also play a role in immune responses. When we spray-inoculated WT, *ttl1ttl3ttl4*, and TTL3-GFP plants (*ttl1ttl3ttl4* complemented line with *TTL3p:TTL3-GFP*, (Amorim-Silva et al., 2019) with the bacterial pathogen *Pseudomonas syringae* pv. tomato DC3000, no differences in bacterial growth were found (Fig. 6A, B). Further, a similar production of apoplastic reactive oxygen species was observed upon flg22 treatment of leaf tissues across the three genotypes (Fig. 6C, D), consistent with the similar formation of the FLAGELLIN SENSING 2 (FLS2)-BRASSINOSTEROID INSENSITIVE 1-ASSOCIATED KINASE 1 (BAK1) complex upon flg22 activation (Fig. 6E). Finally, we determined that mitogen-activated protein kinase (MAPK) phosphorylation is not altered in full seedlings of the *ttl1ttl3ttl4* mutant compared with WT plants (Fig. 6F). Taken together, these results suggest that the BIK1-dependent regulation of TTL proteins is mainly related to cellulose biosynthesis and does not contribute significantly to the plant immune responses tested in this work.

**Fig. 6.**
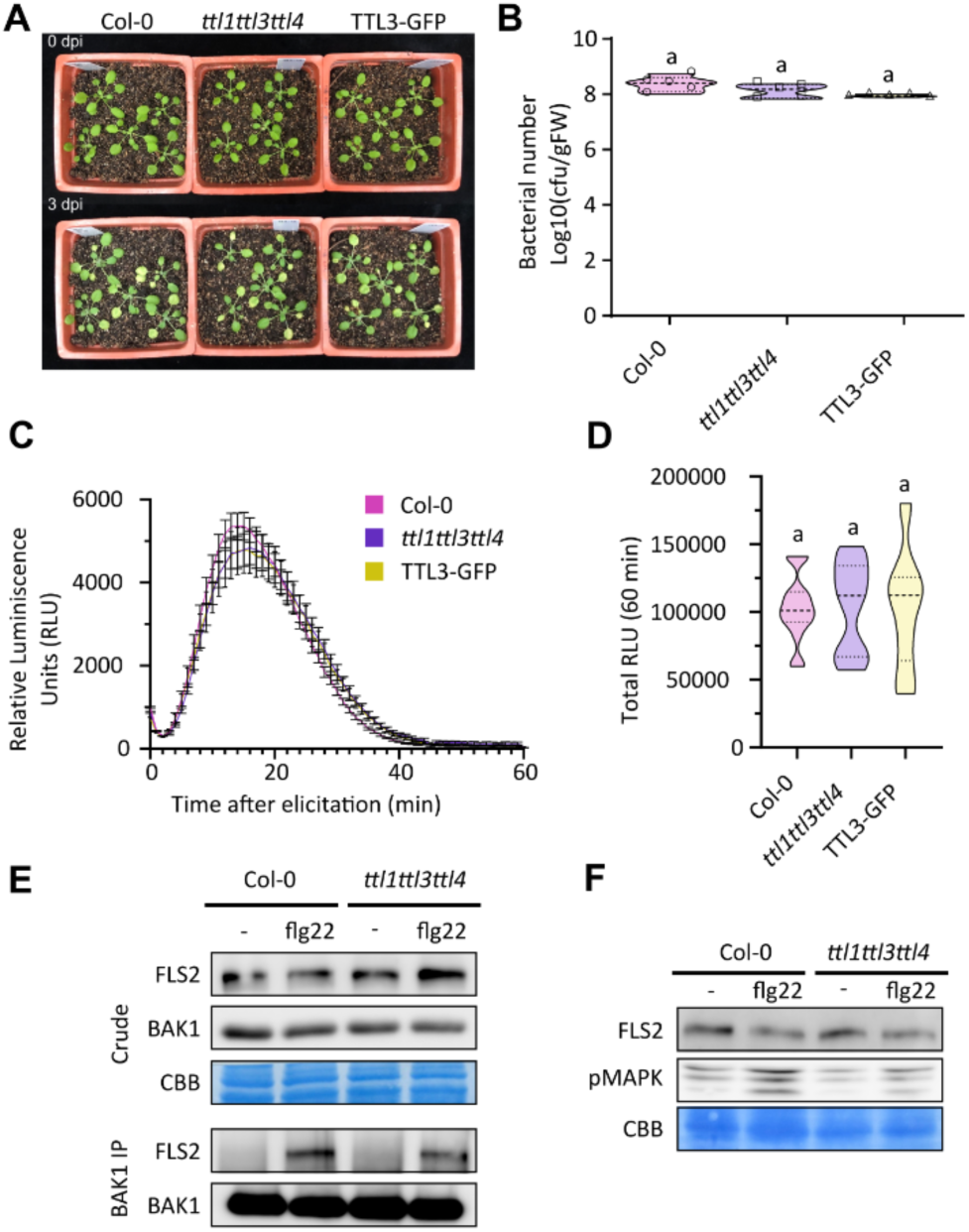
TTL proteins are not involved in immune responses. **A-B.** Three to four-week-old Arabidopsis Col-0, *ttl1ttl3ttl4* triple mutant and TTL3-GFP complemented plants were spray-inoculated with *Pseudomona syringae* pv tomato DC3000. (A) shows representative images of plants before and 3 days post-inoculation (dpi). (B) shows bacterial numbers in seedling tissues 3 dpi, N>5, One way-ANOVA with Tukey’s multiple comparative test and p<0.001, three independent replicates with similar results. **C-D.** Relative (C) and total (D) ROS production was analyzed in those lines upon flg22 treatment through luminescence quantification. One-way ANOVA analysis with Tukey’s multiple comparative test found no difference among genotypes in both assays. **E.** Lack of TTLs in the triple mutant *ttl1ttl3ttl4* does not affect FLS2 or BAK1 accumulation and interaction upon flg22 treatment. **F.** Lack of TTLs in the triple mutant *ttl1ttl3ttl4* does not affect MAPK phosphorylation (activation) in control and flg22 treatment conditions. Equal loading was confirmed by Coomassie Brilliant Blue (CBB) staining of input samples.

## Discussion

This study identifies a phosphorylation-dependent mechanism that links salt-stress perception to the regulation of the CSCs through the coordinated action of TTL3 and BIK1. Our findings establish Ser93 within the N-IDR of TTL3 as the key regulatory residue controlling its subcellular dynamics and identify BIK1 as the kinase responsible for this modification. Together, these results provide a mechanistic framework elucidating how cellulose biosynthesis is stabilized under salt stress.

TTL proteins were previously defined as dynamic CSC-associated components that re-localize from the cytosol to the PM under cellulose-deficient conditions, such as salinity stress (Kesten et al., 2022). Phosphoproteomic analysis identified five phosphorylation sites in the TTL3 N-IDR. A phosphoablative TTL3 variant in these five residues (TTL3^abl^) is constitutively associated with the CSCs, mimicking salt-induced recruitment, whereas a phosphomimetic version (TTL3^mim^) failed to localize properly. Among the five residues, S93 emerged as the primary regulatory residue, as TTL3^S93A^-GFP recapitulated the TTL3^abl^-GFP localization, indicating that phosphorylation at this position is sufficient to control TTL3 PM targeting to associate with CSCs. These results indicate that the level of S93 phosphorylation maintains a cytosolic/CSCs TTL3 ratio under non-stress conditions. At the same time, dephosphorylation promotes CSCs association, indicating that S93 functions as a binary molecular switch that regulates TTL3-CSCs engagement (Fig. 7).

**Fig. 7.**
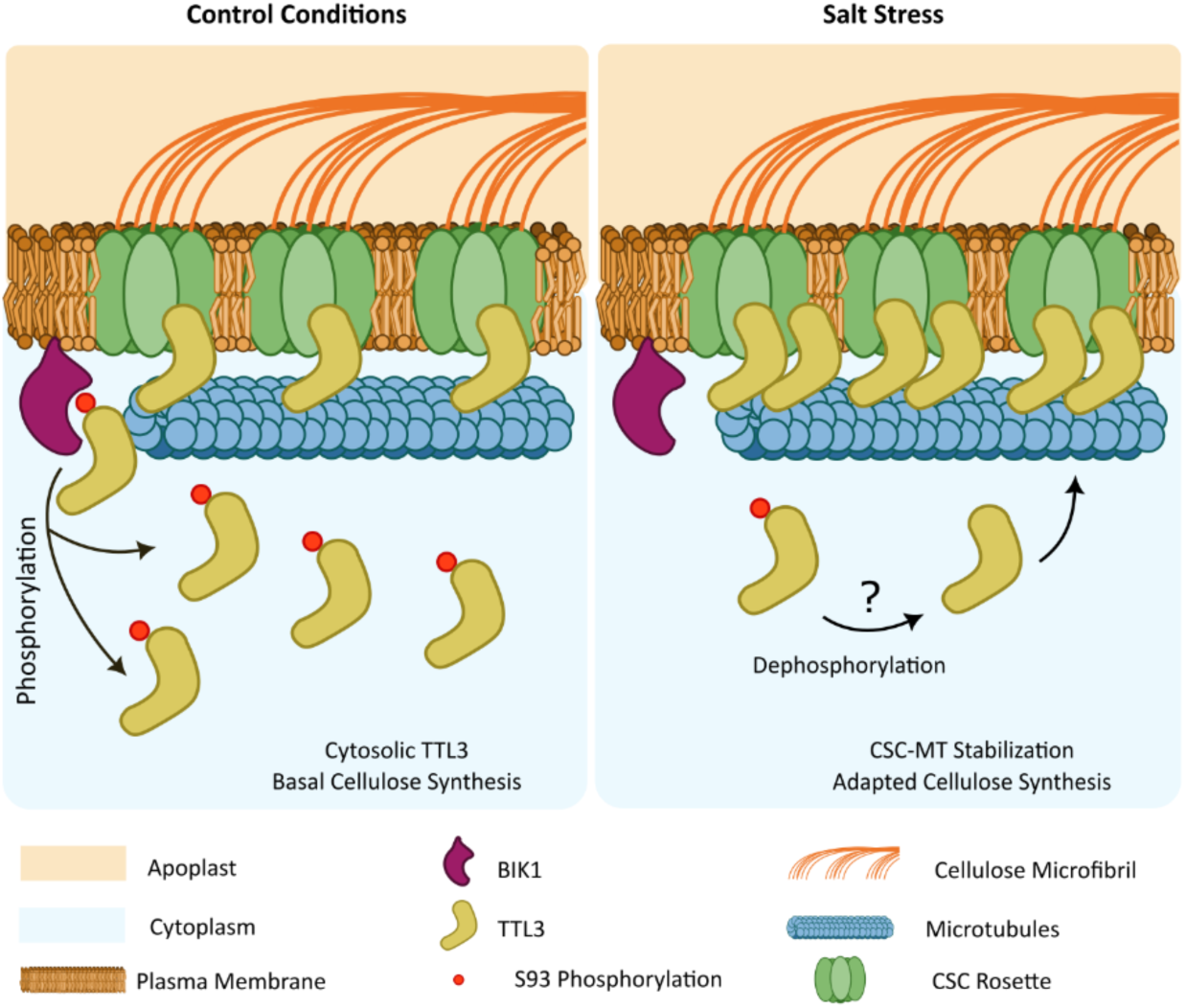
Proposed model for TTL3 regulation of cellulose biosynthesis through BIK1 phosphorylation upon salt stress. Cellulose synthase complexes (CSCs) produce cellulose at plasma membrane under control conditions. In these conditions, TTL3 is partially phosphorylated by BIK1 and confined to the cytosol, while a certain population is unphosphorylated at the plasma membrane, binding the CSCs and the microtubules (MTs). After 28 hours of high salinity, TTL proteins are needed to the formation of a salt-tolerant microtubule network and the CSCs recovery to plasma membrane to restore cellulose synthesis in WT plants. In the absence of TTL proteins, microtubules fail to repolymerize and CSCs fail to repopulate plasma membrane, impairing cellulose biosynthesis. In the case of high salinity, unphosphorylated TTLs either newly synthesized or dephosphorylated by an unknown phosphatase are recruited to the plasma membrane to stabilize CSCs. Phenotypic studies of those lines suggest that lack of phosphorylation and TTL3 accumulation at the CSCs allows cellulose biosynthesis to recover upon high salinity.

IDRs are known to mediate regulatory plasticity through posttranslational modifications (Iakoucheva et al., 2004; Diella et al., 2008). In TTL3, phosphorylation of S93 likely modulates electrostatic properties or conformational flexibility of the N-IDR. Because TTL3 interacts with CESA1 through the IDR (Kesten et al., 2022), phosphorylation at S39 likely inhibits TTL3 access to the CSCs. Phosphorylation of S93 is not only critical for TTL3 localization but also for its function. TTL3^abl^-GFP and TTL3^S93A^-GFP fully complement the root defects of *ttl1ttl3ttl4* as TTL3-GFP, indicating that recruitment of TTL3 to CSCs is essential for sustaining anisotropic expansion during salt stress. These results support a model in which dephosphorylated TTL3 stabilizes CSCs at the PM, thereby preserving cellulose synthesis during stress (Fig. 7).

BIK1 is best characterized for its role in pattern-triggered immunity (Hailemariam et al., 2024; Toth et al., 2026). A recent study has shown that BIK1 phosphorylates SHOU4/SHOU4L, proteins involved in trafficking CESA complexes to the PM, thus establishing a link between cellulose synthesis and pattern-triggered immunity signaling (Wang et al., 2023). Further supporting this convergence, PRR-mediated perception of MAMPs and DAMPs promotes salt tolerance through canonical PTI components, including BIK1 (Loo et al., 2022). Our work extends the function of BIK1 in abiotic stress adaptation by showing that BIK1 phosphorylates the TTL3 N-IDR *in vitro* at S93 and associates with TTL3 via the C-terminal TPR region rather than the N-IDR, indicating spatial separation between the phosphorylation site and the kinase-binding interface. This architecture is consistent with scaffold-like regulation, where binding and phosphorylation are modularly controlled. Interestingly, the C-terminal TPR is also the interaction region for the receptor kinase BRASSINOSTEROID INSENSITIVE 1 (BRI1) in the brassinosteroid signaling pathway (Amorim-Silva et al., 2019), suggesting it is a hub for kinase binding. More importantly, in loss-of-function *BIK1* mutants, the localization of TTL3-GFP mimics that of TTL3^S93A^-GFP, supporting that BIK1 is the main kinase *in vivo* phosphorylating TTL3 S93. As expected, *BIK1* loss-of-function mutants did not show growth defects during salt stress. All these results establish BIK1 as a direct regulator of TTL3 via phosphorylation, controlling its subcellular localization.

Interestingly, a kinase classically associated with immunity directly regulates cellulose biosynthesis during abiotic stress, further highlighting the convergence between cell wall integrity sensing and receptor kinase signaling networks (Wolf, 2022; Zhai et al., 2024). Recently, BIK1 null mutant *bik1-2* has been described to be less responsive to cellulose inhibitor isoxaben (Zhai et al., 2026), increasing the relationship between cell wall integrity and BIK1. This is supported by the unexpected overlapping of DEGs (around 80 %) of a null *BIK1* mutant (*bik1-2*) and the *CESA6* null mutant *prc1-1*. Despite the biochemical connection between TTL3 and BIK1, TTL proteins do not seem to contribute to canonical immune responses measured in leaves or whole seedlings. The triple mutant *ttl1ttl3ttl4* showed normal PTI responses, including *P. syringae* growth, flg22-induced ROS burst, FLS2-BAK1 complex formation, and MAPK activation. This uncoupling suggests that TTLs primarily function in cellulose maintenance under abiotic stress, whereas the roles of BIK1 in immunity appear independent of TTL proteins. However, it cannot be ruled out that TTL proteins might play a role in pathogens whose infection depends on remodeling, loosening, or enzymatic degradation of the host cell wall (Pinto et al., 2025; Carrasco-López et al., 2026).

BIK1 operates as a multifunctional kinase whose downstream targets depend on substrate specificity and cellular context. The TTL-BIK1 module may be a conserved mechanism for the regulation of cellulose biosynthesis in land plants given the conservation of the BIK1 consensus phosphorylated motif present in the TTL family members (Fig. 3A,B), including species in which functional orthologs of BIK1 have been studied, such as Arabidopsis (Paola et al., 2006), tomato (Jaiswal et al., 2022), maize (Li et al., 2022), and *Marchantia polymorpha* (Chu et al., 2023). Overall, this study establishes TTL3 phosphorylation as a crucial node linking RLCK signaling to cellulose biosynthesis under abiotic stress, significantly expanding the functional landscape of BIK1 and deepening our understanding of the maintenance of cell wall integrity.

## Material and methods

### Lines and plant handling

For this work we used *Arabidopsis thaliana* (Col-0 ecotype) and *Nicotiana benthamiana* as plant experimental materials. *Arabidopsis thaliana* T-DNA mutant lines used in this study are *ttl1ttl3* double mutant (Lakhssassi et al., 2012), *ttl1ttl3ttl4* triple mutant (Lakhssassi et al., 2012), and *prc1-1* (AT5G64740, N297) obtained from the European Arabidopsis Stock Centre (NASC: http://arabidopsis.info/). The presence of the T-DNA insert in each mutant was confirmed using dPCR and allele-specific primers.

Transgenic lines *pTTL3:TTL3^abl^-GFP*, *pTTL3:TTL3^mim^-GFP*, *pTTL3:TTL3^S93A^-GFP* were generated by transforming *ttl1ttl3ttl4* with *pTTL3:TTL3^abl^-GFP*, *pTTL3:TTL3^mim^-GFP* or *pTTL3:TTL3^S93A^-GFP*.

Standard procedures and conditions were used for Arabidopsis cultivation. Seeds were surface sterilized using chlorine vapors with a mixture of 100 mL of commercial bleach and 3 mL of 37 % w/w HCl in a sealed container for 4 hours. Afterwards, seed were air-cleared for at least 2 hours in a laminar flow cabinet and plated under sterile conditions on half strength Musharige & Skoog (1/2 MS) pH 5.7 medium supplemented with 1% (w/v) sucrose in control conditions and 0.8% (w/v) plant agar as solidifying agent.

Cellulose biosynthesis altering conditions were established upon the use of 4.5 % or 5.5 % sucrose instead of 1 % used in control plates, supplementing the plates with 100 mM NaCl. In these cases, control plates with 1% of sucrose, no salt was used. For the liquid treatments used for confocal microscopy, the same control media was used and supplemented with 200 mM NaCl when salt stress was used.

Seeds for each genotype used for the same experiment were retrieved from parental plants at the same time to ensure homogeneous germination. Seeds were plated and stratified for 3 days at 4 °C in darkness prior to its growth vertically in a chamber with long-day photoperiod (16 hours light/8 hours darkness), a photon flux of 130±30 mmol photons m^-2^s^-1^ and a temperature of 22 ± 1 °C. Equal germination across genotypes was further confirmed during the elapsing time of the experiment. When required, seedlings were collected and transferred to soil pots with organic substrate and vermiculite (4:1 v/v) and kept under control conditions. Plants were watered every 2 days. For seed collection they were dried and stored under low-humidity conditions and freshly harvested seeds were used for phenotypic analysis.

### Cloning

CRISPR cassettes were generated using GoldenGate cloning method. Shuttle vectors M1 pDGE332 and M2E pDGE334 were used to insert sgRNA sequences in recipient vector pDGE651 containing a Cas9 cassete pRPS5a:Cas9, plant selection of hpt (hygromycin) and FAST, and bacterial selection of Spec, Cm (ccdB). Two sgRNA paired sequences were designed using CHOPCHOP (https://chopchop.cbu.uib.no/). Nucleotides were hybridized mixing 10 µM of each oligo, incubating at 98°C for 5 minutes, and then cooling at RT. A 1:200 dilution of the hybridized oligos were used for the shuttle vectors preparation. 20 fmol of shuttle vector were mixed with 50 fmol of hybridized oligos, ligation buffer, BSA, BpiI and T4 DNA Ligase (0.3 u). Cut/ligation reaction was cycled.

Cut/Ligation reaction was transformed in TOP10 *E. coli* cells and directly used to inoculate a liquid LB culture with carbenicillin. Retrieved plasmids were used for the assembly of plant transformation vectors mixing 20 fmol of the recipient vector, 20 fmol of the shuttle vectors, ligation buffer, BSA, BsaI and T4 DNA ligase (0.3 u). Cut/ligation was cycled as before. The reaction was transformed in *E. coli* for further propagation and then in Agrobacterium for plant transformation.

Constructs for TTL3^abl^-GFP and TTL3^mim^-GFP generation were firstly synthetised by GeneScript, New Jersey, EEUU. We provided them with the first 869 nucleotides of TTL3 CDS sequence which contain the residues that wanted to be swapped to modify alanines or aspartic acids until a SacI restriction site together with the 5’ upstream sequence attL1 gateway sequence containing an ApaI restriction site. Synthetized sequence was retrieved in a pUC57 plasmid which was digested using a SacI and ApaI double digestion at 37 °C for 1 hour and 30 minutes. A pDONR Zeo-TTL3 containing the TTL3 CDS sequence in a Gateway Cloning System suitable vector was also digested with the same enzymes in the same conditions. Both digestion products were run in an agarose electrophoresis gel and mutated sequence and digested WT plasmid without the target region were purified using Gel and PCR Clean-up Nucleospin Kit (Macherey-Nagel, Germany). Purified fragments were mixed and incubated with T4 ligase for 16 hours. Ligation product was transformed in *E. coli* for propagation and then used as pDONR in a LR Multisite Gateway reaction to generate the pENTRY containing the mutated TTL3s driven by its own promoter and fuse to GFP in the C-terminus.

The mutated S93A version of TTL3 were generated using the Q5 Site-directed Mutagenesis Kit of New England Biolabs from pDONR plasmid containing the TTL3 CDS sequence. Primers were designed using the NEB online primer design software https://nebasechanger.neb.com/ flanking the desired nucleotides. PCR amplifications of the plasmid were performed are described in the manufacturer protocol.

Generated linearized plasmid was incubated with the KLD Enzyme Miz at RT for 5’. The mix was transformed in *E. coli* for propagation and the pDONR was used in a Multisite LR Gateway reaction to generate a pENTR in which mutated TTL3 are driven by its own promoter and fused to GFP in the C-terminus. The generated constructs were confirmed by PCR amplification, digestion and sequencing.

### IP-MS

Immunoprecipitation followed by liquid chromatography coupled with tandem mass-spectrometry (IP-LC-MS/MS) was performed to identify TTL3 post-translation modifications as previously described (Wei et al., 2017; Wei et al., 2025), with specific modifications for Arabidopsis tissues.

Five grams of 4-day-old Arabidopsis seedlings were used for large-scale immunoprecipitation of TTL3-GFP. Tissue was frozen in liquid nitrogen and grinded using pestle and mortar. Total proteins were extracted by incubation in ice-cold protein extraction buffer [150 mM Tris-HCl, pH 7.5, 150 mM NaCl, 10% glycerol, 10 mM Ethylene diamine tetra acetic acid (EDTA), 10 mM Dithiothreitol (DTT), 0.5 mM Phenylmethylsulfonyl fluoride (PMSF), 10 mM NaF, 10 mM Na2MoO4, 2 mM NaVO3, 0.5 % (v/v) IGEPAL (IGEPAL CA-630), 1 % (v/v) Plant Protease Inhibitor cocktail (Sigma, St. Louis, MO, USA)] during 40 minutes at 4 °C (from here, the whole procedure was performed inside a cold chamber). Samples were immediately centrifuged for 20 min at 15000 g and 4 °C and supernatants were recovered and filtered through chromatography columns. Equilibrated GFP-trap beads (Chromotek) (100 uL slurry per sample) were added and incubated for 1 hour at 4°C with constant rotation following the manufacturer instructions. Beads were washed 3 times with the same buffer but only 0.2 % detergent and 2 times with no detergent (IGEPAL) and eluted with Laemli buffer. Purification was confirmed by western blot.

TTL3-GFP samples were digested using in-gel digestion by separating proteins through a 10 % SDS-PAGE gel, and the Coomassie Blue-stained bands were excised, reduced by using 10 mM DTT and alkylated with 55 mM iodoacetamide. Samples were digested by trypsin using the standard protocol (Uhrig et al., 2008). Tryptic peptides were extracted using acetonitrile/0.1 % TFA (v/v, 60:40) and dried by refrigerated centrifugal concentrator. Dried peptides were resuspended in 0.1 % v/v formic acid solution and analysed by Q Exactive mass spectrometer (Thermo Electron Finnigan, San Jose, CA). Raw MS/MS data were analyzed using Mascot (v2.5.1) as the search engine, and results were further processed using Scaffold (Proteome Software Inc.). Spectra were searched against an *Arabidopsis thaliana* reference database. Peptide identifications were accepted at greater than 95 % probability, and proteins were retained only if they did not show peptides in the GFP control of 2 independent replicates. Phosphopeptide and phosphosite localization analysis was performed using the Scaffold PTM module.

### Confocal

Confocal images were acquired using a Zeiss LSM880 confocal microscope coupled to an inverted microscope Axio Observer 7. It has a High Sensitivity GaAsP detector, equipped with a 488-nm argon laser for GFP and YFP and 561-nm laser for mC excitation. Images were taken with plan-apochromat 63x/1.4 Oil objective.

### Spinning confocal

*Arabidopsis thaliana* seedlings were imaged at the Center for Advanced Bioimaging of the University of Copenhagen. The microscope consisted of a modular Nikkon Spinning disk confocal with a Revolution XD Imaging System from Andor Microscopy Systems equipped with a 100x oil and 100x resin and laser lines of 488-nm for GFP. For each experiment, two batches of seedlings were prepared for each timepoint and condition with a delay of 25 minutes to ensure imaging timing. At least three seedlings per batch were analyzed, and more than 5 cells per seedling were imaged, with around 30 images for each treatment and timepoint. Microtubules were used for training and ensure focus on plasma membrane

### Skewness

Confocal images were processed using the FIJI Software (Schindelin et al., 2012). Backgrounds were subtracted by the “Substract Background” tool with the sliding paraboloid algorithm with a radius of 50 pixels. The histogram skewness and area of the images was determined using the built-in “Measurement” tool of FIJI. CESA density at plasma membrane was determined using THUNDERStorm plugin (Ovesný et al., 2014). Briefly, images were filtered using a wavelet filter of β-spline, order 3, scale 2.0, the locations of the foci were approximated by the 8-neighbourhood local maximum method, and the subpixel locations of the foci were determined using the Integrated Gaussian point-spread-function with a fitting radius of 3 px and an initial sigma of 1.6 px as described in (Schneider et al., 2022).

### *In vitro* phosphorylation assay

The CDS of the N-terminus of TTL3 was mutated to encode an alanine in the residue 93 as control, transcriptionally fused to Maltose-Binding Protein (MBP) and expressed them in *E. coli*. MBP-MBP was used as negative control. N-terminus of TTL3, TTL3 S93A and negative control were incubated with BIK1, and samples were analyzed through protein electrophoresis. Membrane was stained using Coomassie staining solution, and phosphorylation was detected through the radioactivity of ^32^P.

### Western blotting and co-immunoprecipitation (co-IP)

For western blot analysis of samples, leaves from 3-week-old infiltrated *Nicotiana benthamiana* or 3-day-old Arabidopsis seedlings were collected and grinded to fine powder using liquid nitrogen. Co-IP samples were prepared the same way from the last step of the protocol. Around 100 mg or the Co-IP samples were mixed with the double v/w of 2X Laemmli buffer (125 mM Tris-HCl pH 6.8, 4 % SDS, 20% glycerol, 2 % beta-mercaptoethanol, 0.01 % bromophenol blue). Samples were incubated at 70 °C for 45 minutes and centrifuged at 16,000 g for 5 minutes at RT. Supernatant was used for electrophoresis or kept at -20 °C until use. Electrophoresis was carried out in a 1-mm-width biphasic denaturalizing acrylamide gel with a stacking phase of 5 % acrylamide and pH 6.8 and a resolving phase of 10 % acrylamide pH 8.8. Samples were heated at 70 °C for 20 minutes before loading into the gel dwells, altogether with Spectra™ Broad Range Protein Ladder. Gel ran at 80V until the front reached the resolving gel, and then it was run at 120 V in running buffer (1.92 M glycine, 0.25 M Tris-Base, 1 % SDS). Gel was semi-dry transferred using a Trans-Blot ® Turbo Blotting System into a nitrocellulose membrane activated through 15 s of immersion in methanol and 2 minutes in _dd_H_2_O, and both membrane and Wattman papers were kept in transfer buffer for small and medium proteins (48 mM Tris-Base, 39 mM Glycine, 0.0375 % SDS, 20 % methanol) until electrophoresis was done (around 15 minutes). Transference was performed using the default BioRad Std. transference protocol of the Trans-Blot® Blotting System (constant 0.2 A and maximum of 18V) for 1 h. Transference efficiency was confirmed through gel staining using Coomassie staining solution (0.1 % Brilliant Blue R250, 40 % Ethanol, 10 % glacial acetic acid) for 1 h and distained using distaining solution (40 % ethanol, 10 % glacial acetic acid) for 20 minutes.

Membranes were blocked using 5 % fatty acid free powder milk in TTBS (20 mM Tris-Base, 137 mM NaCl, 0.05 % Tween20) at RT for 2 h. After blocking, membranes were washed using TTBS and incubated on a shaker with the primary antibody at the desired concentration in 1 % fatty acid free powder milk in TTBS at 4 °C O/N. Primary antibody was removed and membrane was washed thrice using TTBS for 10 minutes each. Secondary antibody was added at the desired concentration in 1 % fatty acid free powder milk in TTBS at RT for 1 h. Finally, secondary antibody was removed, and membrane was again washed thrice using TTBS for 10 minutes each. Membranes were revealed using Clarity™ Western ECL Substrate, SuperSignal West Femto™ Maximum Sensitivity or SuperSignal West Atto™ Ultimate Sensitivity in a BioRad Gel Doc XR+ Gel Documentation System for 10 minutes taking images each 10 s unless faster acquisition was needed. Membrane was stained using Coomassie staining solution for 1 h and distaining solution for 20 minutes.

Antibodies used in this work were anti-GFP (1:600, Santa Cruz Technology, sc-9996) andantiHA3 (1:10000, Sigma-Aldrich H3663).

Protein extraction and Co-IP were performed using 500 mg of 3-week-old Nicotiana leaves or 100-300 mg of 3-day-old Arabidopsis seedlings. Samples were frozen and grinded to fine powder in liquid nitrogen. Total protein were extracted with extraction buffer (50 mM Trus-HCl pH 7.5, 150 mM NaCl, 10% glycerol, 10 mM EDTA pH8, 1 mM NaF, 1 mM Na_2_MoO_4_, 10 mM DTT, 0.5 mM PMSF, 1 % v/v P9599 protease inhibitor cocktail [Sigma-Aldrich], Nonidet P-40, CAS:9036-19-5 [USB Amersham Life Science] 0.5 % v/v added at 2 mM/g powder and incubated for 30 minutes at 4°C in an end-over-end rocker. Samples were centrifuged 20 minutes at 4 °C at 9056 g). Supernatants were filtered by gravity through Poly-Prep chromatography columns (731-1550 Bio-Rad), and an aliquot was used as input. The final concentration of detergent (Nonidet P-40) was adjusted to 0.2 % v/v prior to GFP-Trap beads incubation to avoid unspecific binding to the matrix as recommended by the manufacturer. Samples were incubated for 2 h at 4 °C in an end-over-end rocker with 15 µL of ChromoTek GFP-Trap® Magnetic Agarose beads. After incubation, beads were collected with a magnetic rack and washed four times with the wash buffer (same composition as the extraction buffer without detergent). Finally, beads were resuspended in 70 µL of 2X Laemmly sample buffer, heated and analyzed through protein electrophoresis and western blot as described in “Protein electrophoresis and western blot analysis”.

### RNAseq

For RNAseq analysis, pools of 12, 4-week-old Arabidopsis rosettes, were powdered in liquid N2 and total RNA was isolated using the E.Z.N.A.® Plant RNA Kit (Omega Bio-Tek). RNA integrity was checked by visualization in a 2 % agarose gel electrophoresis. For the library preparations and sequencing on an Illumina Novaseq platform the samples were sent to the Novogene (UK) Company Ltd, and 150-bp paired-end reads were generated. The reads were quality filtered using fastp 1.0 software (Chen, 2025). The resulting reads were then aligned to the TAIR10.1 version of the *Arabidopsis thaliana* genome sequence (https://www.arabidopsis.org/) using Hisat2 version 2.2.1 (Kim et al., 2015). The resulting read alignments were visualized using Integrative Genomics Viewer software (Robinson et al., 2011) for validation of each genotype. These read alignments (in BAM format) were used for transcript quantification with the featureCounts 2.0.6 program (Liao et al., 2014). To determine DEGs, a False Discovery Rate (q-value) cutoff of ≤ 0,05 was set. DEGs were subjected to singular enrichment analysis for the identification of overrepresented GO terms using ShinyGO tool (https://bioinformatics.sdstate.edu/go/; (Ge et al., 2020)) with the default options.

### Immune assays

Bacterial inoculation assays were performed as previously described (Wang et al., 2019). *Pseudomonas syringae pv tomato* DC3000 (*Pto* DC3000) was streaked on selective ½ salt LB plates (10 g Tryptone, 5 g Yeast Extract, 5 g NaCl, 15 g Agar per 1 L) and cultivated at 22 °C for 2 d. Bacteria were collected from the plates into sterile water, the OD was adjusted to a value of 0.1 (5x10^7^ cfu/mL) or 0.2 (10^8^ cfu/mL), and silwet-L77 was added to a final concentration of 0.03% before performing spray inoculation onto 3- to 4-week-old soil-grown Arabidopsis under short day conditions. The plants were then covered with cling wrap for 3 h. The whole plants were harvested in 1.5-mL microcentrifuge tubes and weighed. Plant tissues were ground and homogenized in sterile water before plating serial dilutions on selective ½ salt LB agar plates. The plates were placed at 28 °C for 1.5 d and the bacterial growth was calculated as colony-forming units.

ROS burst assays were conducted as previously described (Sang and Macho, 2017). Briefly, 4-mm leaf discs from 5- to 6-week-old Arabidopsis plants grown in short day conditions were transferred to 96-well microplates (PerkinElmer, Waltham, MA, US) with 100 μl Mili-Q water per well and incubated overnight. Water was then removed and ROS burst was elicited by adding 100 μl of a solution containing 100 ng flg22, 100 μM luminol, and 20 μg/mL horseradish peroxidase. The luminescence was recorded over 40 minutes using a Thermo Scientific VARIOSKAN FLASH (Thermo Fisher Scientific, Waltham, MA, US).

MAPK activation assays were performed as previously described (Macho et al., 2012). Briefly, 12-day-old seedlings (5 d on 1/2 MS solid agar plates, then 7 d in 1/2 MS liquid medium) were treated with a 100 nM flg22^Pto^ elicitor solution (or water as mock treatment). Samples (2 seedlings per treatment) were collected 15 minutes after treatment. Proteins were extracted and subjected to immunoblots with anti-pMAPK antibody (Cell Signaling, 4370S).

### Statistics and graphs

Statistical analyses were conducted using Prism 8.0.2 for Microsoft (GraphPad Software, www.graphpad.com). Used test include unpaired t-test, one-way or two-ways analysis of variance (ANOVA) follower by Tukey’s multiple comparison test or Dunnett’s multiple comparisons test, respectively. P value < 0.05 if not stated otherwise.

Graphs were performed in Prism 8.0.2 for Microsoft (GraphPad Software, www.graphpad.com).

## Author contributions

Conceptualization: P. F., P.-M. R., A.-S. V., B. M. A.

Data curation: P. F., P.-M. R.

Formal analysis: P. F., P.-M. R.

Funding acquisition: P. F., P.-M. R., P. S., M. A. P., P.-S. J., Z. C., DF. T., R. L., A.-S. V., B. M. A.

Investigation: P. F., P.-M. R., D. E. A., L. J., T. R., DF. T.

Methodology: P. F., P.-M. R., D. E. A., C. L.

Project administration: P. F., P.-M. R., A.-S. V., B. M. A.

Supervision: M. A. P., Z. J.-M., Z. C., P. S., A.-S. V., B. M. A.

Validation: P. F., P.-M. R.

Visualization: P. F., P.-M. R., D. E. A., A.-S. V.

Writing – original draft: P. F., P.-M. R., A.-S. V., B. M. A.

Writing – review & editing: P. F., P.-M. R., D. E. A., M. A. P., P.-S. J., T. R., DF. T. A., Z. J.-M., Z. C., R. L., P. S., A.-S. V., B. M. A.

## Acknowledgments

F.P. acknowledges Jose Enrique Perez Martin from the University of Malaga (ORCID: 0009-0003-8039-1944) for his contribution to data analysis automation in R.

## Tables

Tables S1, S2 and S3 are uploaded to the website.

---

**Sup. Fig S1.**
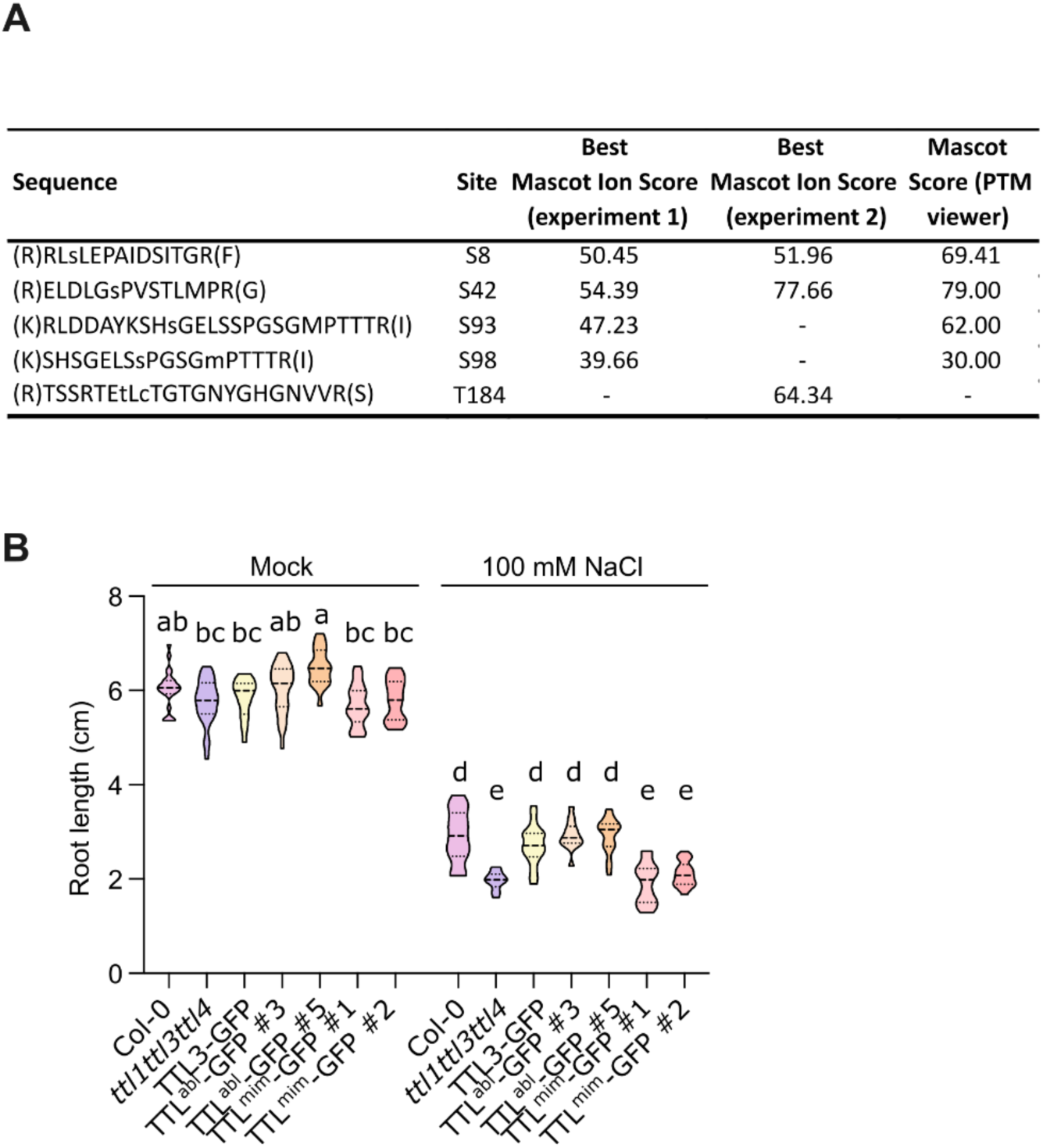
Raw data for phosphorylation studies of TTL3 IDR. **A.** IP-MS PTM analysis of TTL3. Residues within the N-IDR and Mascot Ion Score higher than 35 were considered for each replicate. **B**. Raw data for root length quantifications of relative root length shown in Fig. 1F. Two way-ANOVA with Tukey’s multiple comparative test and p<0.001. Three independent replicates with similar results.

**Sup. Fig. S2:**
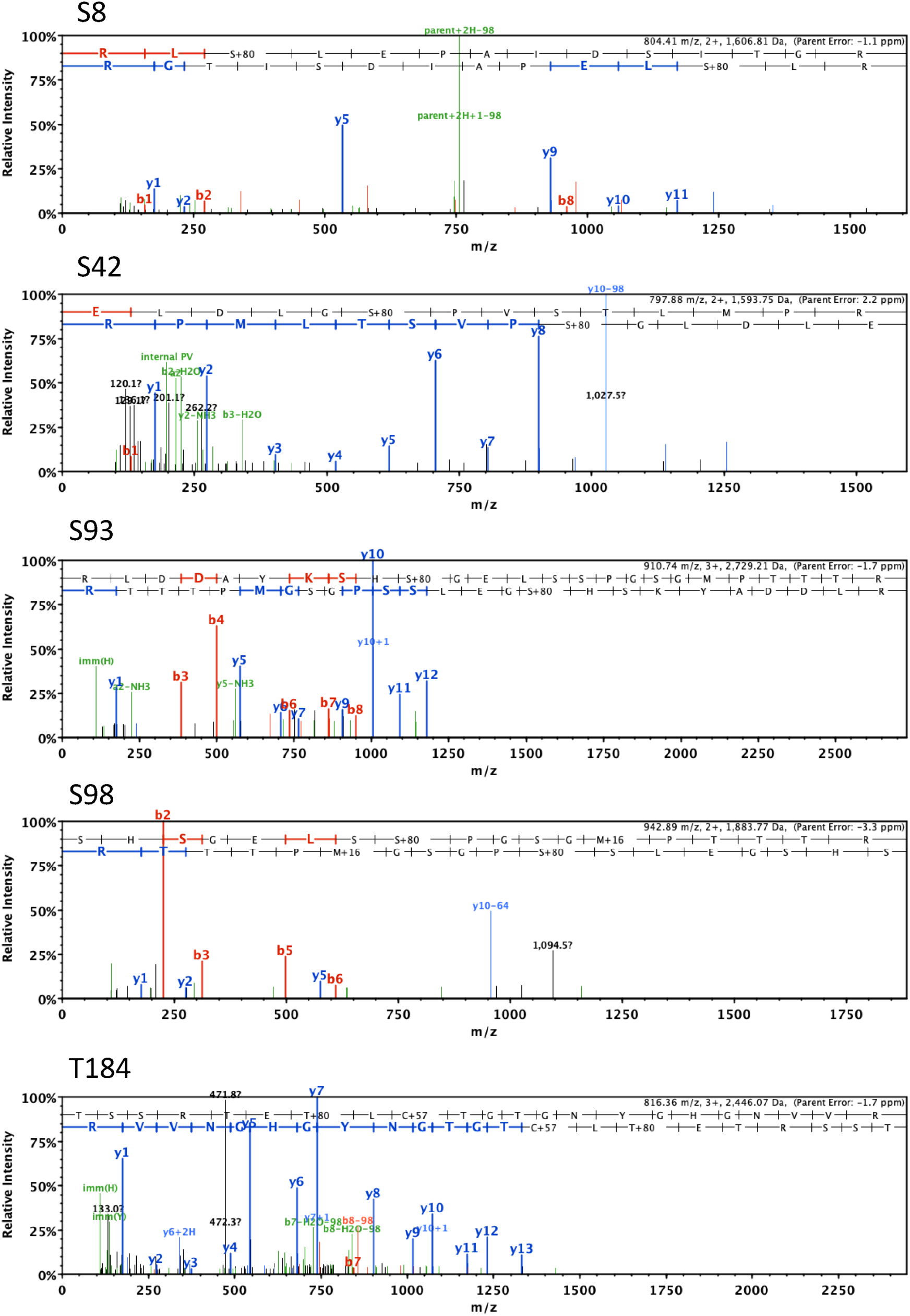
Representative mass spectra for the phosphorylated peptides described in Figure S1.

**Sup. Fig. S3.**
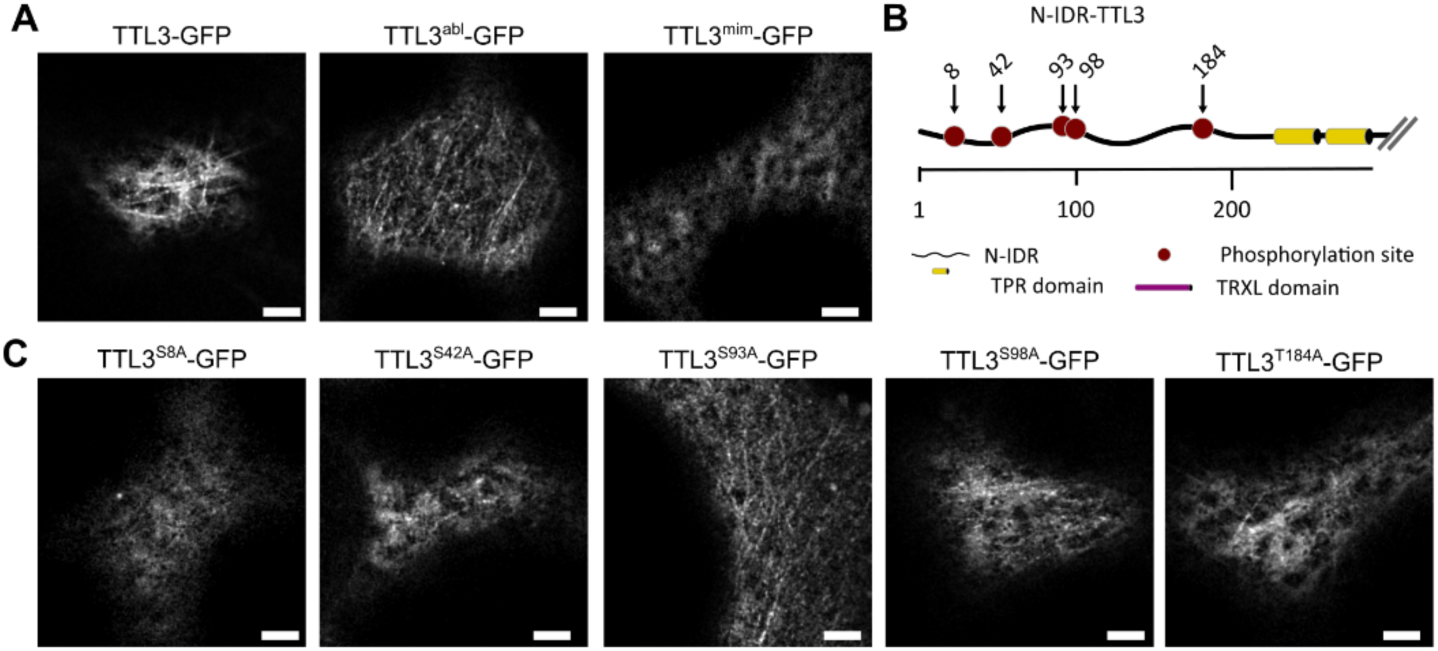
*Nicotiana benthamiana* transient expression screening of mutated TTL3. **A.** Representative image of TTL3-GFP, TTL3^abl^-GFP and TTL3^mim^-GFP as screening controls transiently expressed in *Nicotiana benthamiana* leaves. **B.** Schematic representation of the N-IDR of TTL3 depicting the mutated residues in the screening. **C.** Representative image of individually serine/threonine-mutated TTL3-GFP at the residues 8, 42, 93, 98, 184 to alanine transiently expressed in *Nicotiana benthamiana* leaves. Scale bars 10 μm.

**Sup. Fig. S4.**
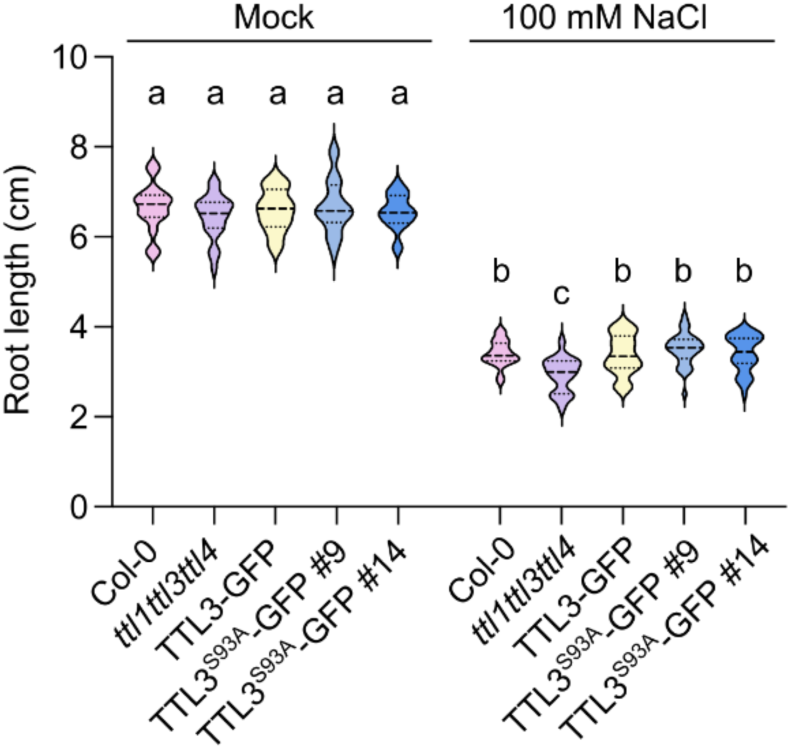
Raw data for root length quantifications of relative root length shown in Fig. 2C. Two way-ANOVA with Tukey’s multiple comparative test and p<0.001. Three independent replicates with similar results.

**Sup. Fig. S5.**
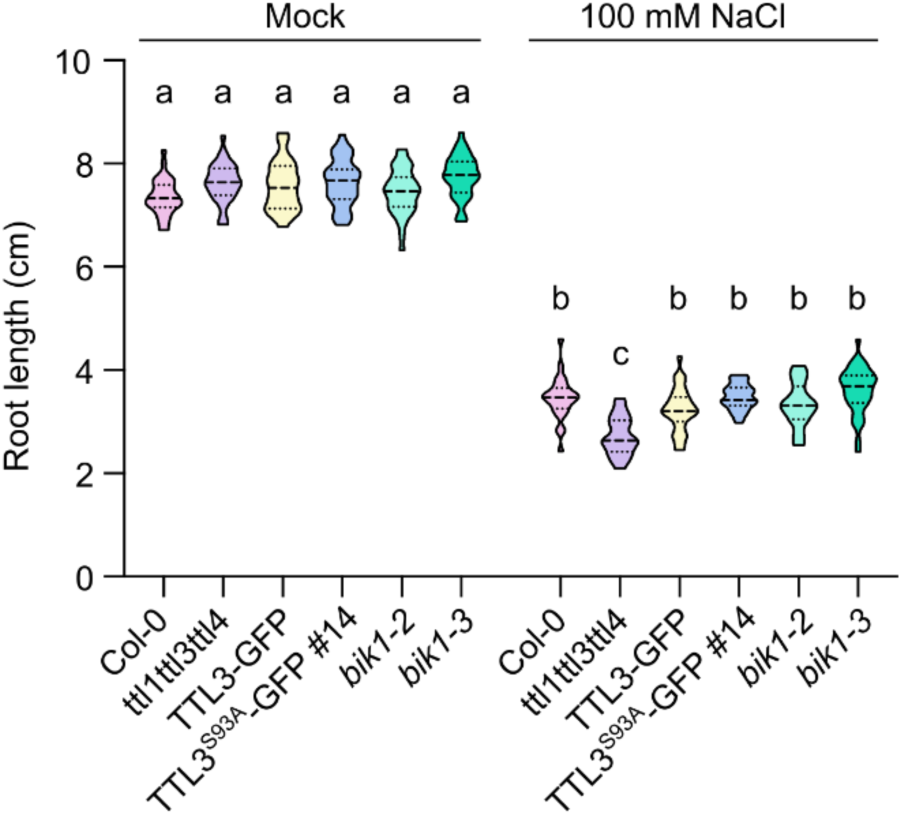
Raw data for root length quantifications of relative root length shown in Fig. 4E. Data from Fig 4F. Two way-ANOVA with Tukey’s multiple comparative test and p<0.001. Three independent replicates with similar results.

**Sup. Fig. S6.**
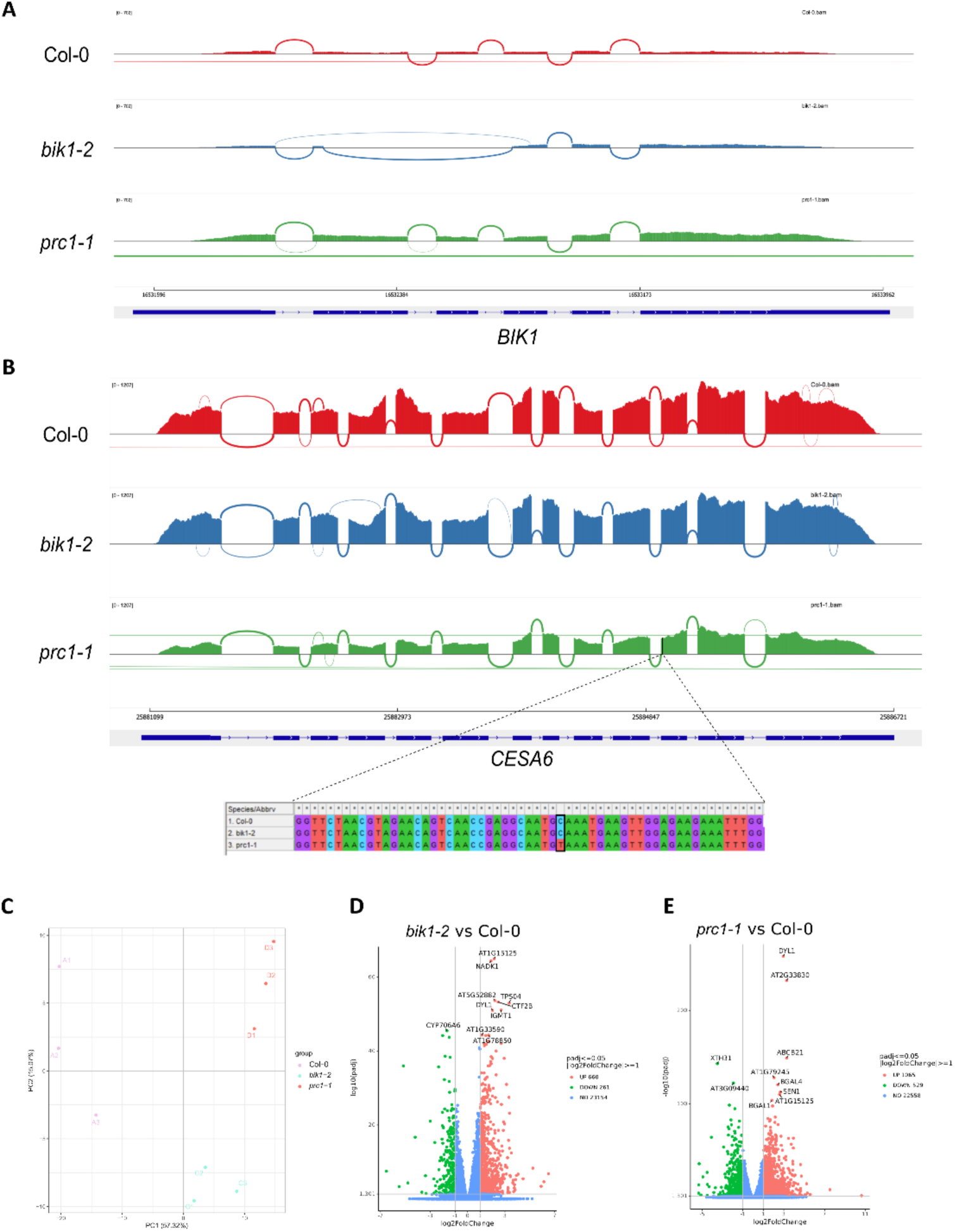
Transcriptome profiles of four-week-old Arabidopsis rosettes. **A-B.** RNAseq mapped reads showing the *bik1-2* mutation in the *BIK1* gene and the *prc1-1* mutation (C2159T, Q720stop) in the *CESA6* gene visualized using Integrative Genomics Viewer (IGV). **C.** Principal Component Analysis plot displaying all replicates for each of the three genotypes. **D-E.** Volcano plots showing that the statistically significant DEGs (red-up and green-down). Fold change between each mutant and Col-0 (WT) is presented in log2 scale, whereas adjusted p-value (padj) are presented in -log10 scale. Significance cutoff was set at α=0.05.

